# Global Adoption of High-Sensitivity Cardiac Troponins and the Universal Definition of Myocardial Infarction

**DOI:** 10.1101/371138

**Authors:** Atul Anand, Anoop SV Shah, Agim Beshiri, Allan S Jaffe, Nicholas L Mills

## Abstract

**Importance:** The third Universal Definition of Myocardial Infarction aimed to standardize the approach to the diagnosis and management of myocardial infarction. High-sensitivity cardiac troponin testing was recommended, as these assays have improved precision at low concentrations, but concerns over specificity may have limited implementation.

**Objective:** To determine the global adoption of high-sensitivity cardiac troponin assays and key recommendations from the Universal Definition.

**Design, Setting and Participants:** Global survey of 1,902 medical centers across 23 countries evenly distributed across all five continents. Included respondents were involved in the diagnosis and management of patients with suspected acute coronary syndrome at their institutions.

**Main Outcomes and Measures:** Structured questionnaire detailing the primary biomarker used for myocardial infarction, diagnostic thresholds and critical elements of clinical pathways for comparison to the third Universal Definition recommendations.

**Results:** Cardiac troponin was the primary diagnostic biomarker for myocardial infarction at 96% of all sites surveyed. Only 41% of centers had adopted high-sensitivity cardiac troponin assays, with wide variation from 7% in North America to 60% in Europe. Sites using high-sensitivity assays more frequently employed serial sampling pathways (91% *vs.* 78%) and the 99^th^ percentile diagnostic threshold (74% *vs.* 66%) when compared to sites using the previous generation of troponin assays. Furthermore, sites using high-sensitivity assays more often used earlier serial sampling (≤3 hours) and accelerated diagnostic pathways. However, fewer than 1 in 5 sites using high-sensitivity assays had adopted sex-specific thresholds (18%).

**Conclusions and Relevance:** Progress has been made in adopting the recommendations of the Universal Definition of Myocardial Infarction, particularly in the use of the 99^th^ percentile diagnostic threshold and serial sampling. However, high-sensitivity assays are used in a minority of sites and sex-specific thresholds in even fewer. These findings highlight regions where additional efforts are required to improve the risk stratification and diagnosis of patients with myocardial infarction.

## Introduction

In 2012, a Joint ESC/ACCF/AHA/WHF Task Force presented the third Universal Definition of Myocardial Infarction (UDMI).^1^ This landmark document continued efforts to standardize the classification of a condition that affects 790,000 patients each year in the United States alone.^2^ The following key recommendations were made. Cardiac troponin is the preferred biomarker of myocardial injury due to its high specificity and sensitivity. Serial sampling should be performed at presentation and after 3-6 hours to demonstrate a rise and/or fall in troponin, except in those presenting late with a high pre-test probability of myocardial infarction. The diagnostic threshold for myocardial infarction is defined by the 99^th^ percentile upper reference limit (URL) of cardiac troponin in a healthy reference population. The task force emphasized analytical precision and recommend use of high-sensitivity cardiac troponin assays capable of <10% coefficient of variation at the diagnostic threshold. Finally, it was recommended that sex-specific thresholds be adopted when using high-sensitivity assays, where there is evidence of variation between healthy males and females in reference populations.

Since publication of the third UDMI, high-sensitivity cardiac troponin use has increased worldwide, but these assays have only recently been approved for use in the United States. Accordingly, we assessed the adoption of these guideline recommendations through a comprehensive global survey.

## Methods

A structured telephone survey was completed with staff from the laboratory, emergency or cardiology departments of 1,902 clinical institutions across 23 countries between February and April 2016. Included sites provided balanced sampling from five regions: North America (United States and Canada), Europe (France, Germany, Italy, Spain and United Kingdom), Latin America (Argentina, Brazil and Mexico), Asia Pacific (Australia, China, India and Japan) and the Middle East and Africa (Egypt, Kenya, Morocco, Saudi Arabia, South Africa, Tanzania, Turkey, Uganda and United Arab Emirates). Data was only collected from respondents who confirmed knowledge of guidelines for the diagnosis and assessment of myocardial infarction at their institution, including laboratory biomarkers tested. The structured questionnaire recorded the sensitivity of biomarkers, thresholds for myocardial infarction and diagnostic pathway for each institution (**see Supplement**).

## Results

Cardiac troponin was the primary diagnostic biomarker for myocardial infarction in 96% of surveyed centers. Outside of Latin America, CKMB use was very limited (**Table**). Significant global variation was observed in the adoption of high-sensitivity cardiac troponin assays, from 7% in North America to 60% in Europe (**Figure 1A and B**). Overall, a minority (41%) of surveyed sites had adopted high-sensitivity assays. Sites using high-sensitivity assays more frequently utilised serial sampling strategies (91% *vs.* 78%) and the 99^th^ percentile diagnostic threshold (74% *vs.* 66%) when compared to sites using contemporary assays (**Figure 1C**).

**Table:** Primary diagnostic assay and assessment of UDMI criteria amongst high-sensitivity cardiac troponin users by country. ^a^Africa = Kenya, Tanzania and Uganda (results considered together). cTn = cardiac troponin; CK-MB = creatine kinase-MB; UDMI-Universal Definition of Myocardial Infarction. For UDMI criteria, percentage per country calculated as a proportion of those sites providing a definitive response to each criterion.

| Country | Number of sites | Primary diagnostic assay |  |  | UDMI Criteria for high-sensitivity sites (n=774) |  |  |  |
| --- | --- | --- | --- | --- | --- | --- | --- | --- |
|  |  | High-sensitivity cTn, n (%) | Contemporary sensitivity cTn, n (%) | CK-MB, n (%) | 99<sup>th</sup> percentile threshold, n (%) | Serial sampling, n (%) | Serial sample at 3-6 hours, n (%) | Sex-specific thresholds, n (%) |
| United Arab Emirates | 50 | 39 (78) | 11 (22) | 0 (0) | 13/15 (87) | 38/38 (100) | 17/31 (55) | 2/36 (6) |
| United Kingdom | 80 | 58 (72) | 22 (28) | 0 (0) | 34/43 (79) | 56/57 (98) | 42/54 (78) | 22/58 (38) |
| Italy | 82 | 55 (67) | 26 (32) | 1 (1) | 29/47 (62) | 54/55 (98) | 49/54 (91) | 21/55 (38) |
| Saudi Arabia | 67 | 42 (63) | 21 (31) | 4 (6) | 16/23 (70) | 38/39 (97) | 18/33 (55) | 9/42 (21) |
| Africa<sup>a</sup> | 10 | 6 (60) | 4 (40) | 0 (0) | 0/1 (0) | 6/6 (100) | 5/5 (100) | 0/6 (0) |
| France | 80 | 46 (57) | 34 (42) | 0 (0) | 29/41 (71) | 45/46 (98) | 37/44 (84) | 10/46 (22) |
| Australia | 100 | 55 (55) | 45 (45) | 0 (0) | 45/46 (98) | 53/55 (96) | 40/47 (85) | 11/54 (20) |
| Spain | 80 | 44 (55) | 36 (45) | 0 (0) | 32/37 (86) | 42/43 (98) | 31/41 (76) | 5/44 (11) |
| India | 100 | 53 (53) | 45 (45) | 2 (2) | 43/48 (90) | 36/46 (78) | 32/35 (91) | 4/52 (8) |
| Argentina | 100 | 52 (52) | 29 (29) | 19 (19) | 21/31 (68) | 46/51 (90) | 32/43 (74) | 13/52 (25) |
| Germany | 80 | 40 (50) | 37 (46) | 3 (4) | 27/33 (82) | 33/39 (85) | 23/30 (77) | 7/39 (18) |
| Japan | 100 | 49 (49) | 48 (48) | 3 (3) | 22/38 (58) | 26/44 (59) | 11/17 (65) | 2/48 (4) |
| South Africa | 121 | 58 (48) | 61 (50) | 2 (2) | 33/41 (80) | 53/56 (95) | 38/45 (55) | 2/58 (3) |
| Brazil | 100 | 37 (37) | 61 (61) | 2 (2) | 10/29 (34) | 35/37 (95) | 18/22 (82) | 3/32 (9) |
| Turkey | 122 | 45 (37) | 74 (61) | 3 (2) | 17/26 (65) | 35/37 (95) | 13/24 (54) | 9/44 (20) |
| Morocco | 23 | 8 (35) | 14 (61) | 1 (4) | 3/6 (50) | 5/8 (63) | 4/5 (80) | 2/8 (25) |
| China | 100 | 34 (34) | 62 (62) | 4 (4) | 31/34 (91) | 30/31 (97) | 12/15 (80) | 7/34 (21) |
| Canada | 100 | 27 (27) | 73 (73) | 0 (0) | 7/17 (41) | 23/27 (85) | 20/22 (91) | 3/27 (11) |
| Mexico | 100 | 26 (26) | 37 (37) | 37 (37) | 10/17 (59) | 18/25 (72) | 14/16 (88) | 6/26 (23) |
| Egypt | 7 | 0 (0) | 7 (100) | 0 (0) | – | – | – | – |
| United States of America | 300 | 0 (0) | 297 (99) | 3 (1) | – | – | – | – |
| **Total, n (%)** | **1902** | **774 (41)** | **1,044 (55)** | **84 (4)** | **422/573 (74)** | **672/740 (91)** | **456/583 (78)** | **138/761 (18)** |

**Figure 1:**
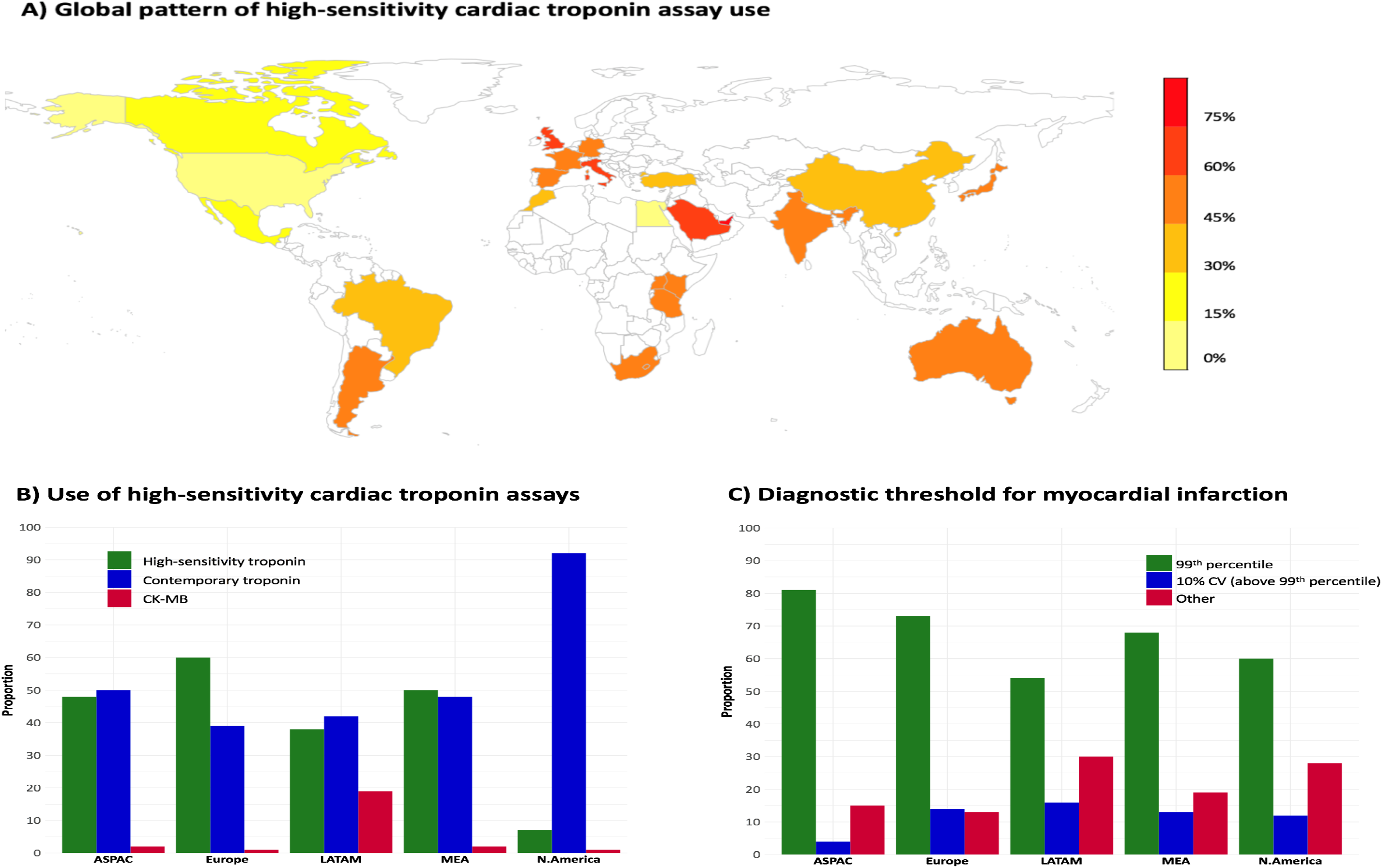
Global adoption of high-sensitivity cardiac troponin assays and diagnostic thresholds for myocardial infarction. ASPAC = Asia-Pacific (Australia, China, India and Japan); Europe (France, Germany, Italy, Spain and United Kingdom);
LATAM = Latin America (Argentina, Brazil and Mexico); MEA = Middle East and Africa (Egypt, Kenya, Morocco, Saudi
Arabia, South Africa, Tanzania, Turkey, Uganda and United Arab Emirates); N.America = North America (United States
and Canada)

Serial sampling 3-6 hours after symptom onset was reported by 78% of high-sensitivity sites. There was a shift towards earlier serial sampling at ≤3 hours in these institutions (52% *vs.* 35% of contemporary users, **Figure 2**). However, diagnostic pathways including sex-specific thresholds were only reported by a minority (18%) of high-sensitivity sites (**Table**).

**Figure 2:**
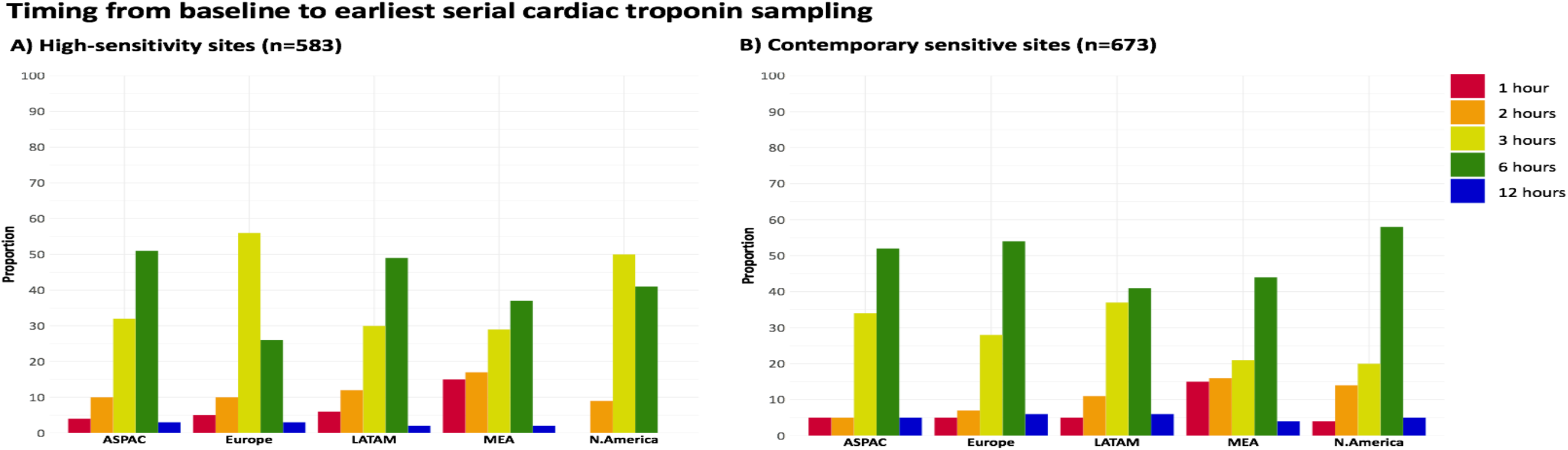
Comparison of serial sampling patterns for high-sensitivity and contemporary sensitive cardiac troponin sites. ASPAC = Asia-Pacific (Australia, China, India and Japan); Europe (France, Germany, Italy, Spain and United Kingdom);
LATAM = Latin America (Argentina, Brazil and Mexico); MEA = Middle East and Africa (Egypt, Kenya, Morocco, Saudi
Arabia, South Africa, Tanzania, Turkey, Uganda and United Arab Emirates); N.America = North America (United States
and Canada)

## Discussion

Five years after publication of the third UDMI, this survey of almost 2,000 institutions across 5 continents addresses progress towards the aims of the Global Task Force and makes a number of important observations. First, cardiac troponin has become ubiquitous as the diagnostic biomarker of myocardial infarction, although CK-MB usage persists in some parts of Latin America. Second, although the majority of centers have adopted the 99^th^ percentile URL and serial testing at 3-6 hours after symptom onset, only a minority are using high-sensitivity assays that provide increased precision and improved information at this threshold. Thirdly, few sites using high-sensitivity assays reported sex-specific diagnostic thresholds.

As recently as 2001, the Euro Heart study determined fewer than two thirds of patients with suspected acute coronary syndrome were assessed with cardiac troponin, often in combination with a second biomarker.^3^ Within two decades, we have observed a rapid shift to near universal use of cardiac troponin. There is also clear evidence of movement towards lower diagnostic thresholds defined by healthy reference populations rather than limits of assay precision. The CARMAGUE survey in 2006 reported only 28% of troponin T and 42% of troponin I sites in Europe were using the 99^th^ percentile URL.^4^ This had risen to 52% across both assays in a 2013 survey by the same group.^5^ In this global study, use of the 99^th^ percentile URL has risen to 70% across all sites whether using a high-sensitivity or previous generation assay.

However, the clinical impact of lower diagnostic thresholds remains contentious, particularly in the era of high-sensitivity troponin assays. The potential benefits of increased sensitivity may be partially mitigated by reduced specificity for type 1 myocardial infarction with more frequent identification of type 2 events secondary to supply/demand imbalance and detection of myocardial injury.^6^ However, this may be because many centers have previously not used the 99^th^ percentile URL even with contemporary cardiac assays.^5,7^ Increased recognition of type 2 myocardial infarction may not advance patient outcomes and formal treatment trials are lacking. However, lower diagnostic thresholds improve overall outcomes for suspected acute coronary syndrome populations.^8^ It is therefore encouraging that 74% of high-sensitivity and 66% of contemporary troponin users report adoption of the recommended 99^th^ percentile threshold.

One approach to improve specificity is to assess dynamic changes in cardiac troponin concentration with early repeat testing. Although the majority of sites follow the current UDMI criteria for serial sampling at presentation and 3-6 hours, it is clear those with access to high-sensitivity assays are increasingly performing rapid retesting at 1 and 2 hours from presentation to facilitate earlier hospital discharge. Numerous strategies have been tested and some even recommended within regional guidelines, but controversy remains about optimum deployment of these early rule-out approaches.^9-12^ The next UDMI update offers an opportunity to better harmonize the approach to the safe use of early rule-out thresholds below the 99^th^ percentile, analogous to the third UDMI’s standardization of the diagnostic threshold for myocardial infarction.

Global variation in the selection of assay and diagnostic thresholds impacts on the reporting of myocardial infarction events in multi-centered international trials that inform guidelines. This has been demonstrated across the 276 hospital laboratories contributing to the ISCHEMIA trial, where 1 in 4 sites reported decision thresholds more than 5-fold greater than manufacturer recommendations, with the majority using contemporary cardiac troponin assays.^13^ It is clear that variation also exists in the adoption of sex-specific diagnostic thresholds, despite evidence of potential benefit to reduce the under-recognition of myocardial infarction in women.^14,15^ This may reflect dominance of the high-sensitivity cardiac troponin T assay which, until its U.S. approval, has not recommended sex-specific cut-off values.

Our global survey does have some limitations. Not all countries could be included, but as the largest such study and the first to report across every continent, extensive efforts were made to provide representative sampling. We have taken a truly global perspective, including many developing countries. It is critical to continue standardization towards a global consensus on both the identification and safe rule-out of myocardial infarction. This will improve the efficiency and efficacy of healthcare systems and permit comparisons between countries and regions to further enhance standards of care. Efforts are still required to ensure all countries have access to high-sensitivity troponin assays and report using the guideline recommended thresholds.

## Acknowledgments and funding

This study was designed by ASVS, AB and NLM. AA was responsible for data analysis. AA, ASVS and NLM drafted the initial manuscript that was approved by all authors. The survey was funded by Abbott Diagnostics and was conducted by an independent organisation, Suazio Consulting. Abbott Diagnostics provided no study funding. AA is supported by a research fellowship from Chest Heart and Stroke Scotland (15/A163) and NLM by a British Heart Foundation Butler Senior Clinical Research Fellowship (FS/16/14/32023) and Special Project Grant (SP/12/10/29922).

